# Human Epigenetic Aging is Logarithmic with Time across the Entire LifeSpan

**DOI:** 10.1101/401992

**Authors:** Sagi Snir, Matteo Pellegrini

**Affiliations:** Dept. of Evolutionary Biology, University of Haifa, Israel; Dept. of Molecular, Cell and Developmental Biology; University of California, Los Angeles, CA 90095, USA

## Abstract

It is well established that organisms undergo epigenetic changes both during development and aging. Developmental changes have been extensively studied to characterize the differentiation of stem cells into diverse lineages. Epigenetic changes during aging have been characterized by multiple epigenetic clocks, that allow the prediction of chronological age based on methylation status. Despite their accuracy and utility, epigenetic age biomarkers leave many questions about epigenetic aging unanswered. Specifically, they do not permit the unbiased characterization of non-linear epigenetic aging trends across entire life spans, a critical question underlying this field of research. Here we a provide an integrated framework to address this question. Our model, inspired from evolutionary models, is able to account for acceleration/deceleration in epigenetic changes by fitting an individuals model age, the *epigenetic age*, which is related to chronological age in a non-linear fashion. We have devised a two stage procedure leveraging these model ages to infer aging trends over the entire lifespan of a population. Application of this procedure to real data measured across broad age ranges, from before birth to old age, and from two tissue types, suggests a universal logarithmic trend characterizes epigenetic aging across entire lifespans. This observation may have important implications for the development and application of future, more accurate, aging biomarkers.

## 1 Introduction

Cell type specific differences in gene expression are partially controlled by chromatin accessibility and specific covalent modifications. These include modification to histones and DNA [25, 2]. Among these, DNA methylation has been one of the most extensively studied components of cell type specification [11, 5]. The covalent attachment of a methyl group to cytosine is catalyzed by either de novo or maintenance methyltransferases, and in mammals is primarily targeted to CpG dinucleotides. Most CpGs in mammalian genomes are methylated, but pockets of hypomethylation exist, largely at promoters and enhancers. It has been shown that the absence of DNA methylation is closely associated with the presence of H3K4 methylation, which is also a hallmark of enhancers and promoters [16]. As stem cells differentiate along myriad lineages, each cell type tends to have distinctive DNA methylation profiles, mostly due to differential activation of enhancers [8].

Once an organism reaches its adult stage, these cell types and their respective epigenomes, have been largely determined. While these developmental epigenetic changes are believed to be rapid and extensive, in the past few years it has become ever more apparent that epigenetic changes continue to occur as an organism ages [21]. This observation has led to the development of multiple epigenetic clocks, that is, biomarkers that accurately predict the chronological age of an individual based on his/her DNA methylation profile [9, 7]. These epigenetic clocks have been extensively used in aging research and have proven to be more accurate than previous aging biomarkers, such as the length of telomeres [14]. Using these epigenetic clocks, much has been learned about the effects of the environment on aging. For example, it is well known that the restriction of calories in mice slows down aging, increases lifespan as well as the rate of the epigenetic clock [23]. Similar conclusions have been found in humans, where individuals with more rapid epigenetic aging tend to suffer from higher all-cause mortality rates [4].

While these epigenetic clocks have proven to be useful for aging research, they are constructed using machine learning methods that provide limited insights into the underlying processes that are driving these changes. For example, the Horvath epigenetic clock biomarker is constructed by selecting 353 CpG sites using penalized lasso regression, that optimally predict the chronological age of an individual. This biomarker also sets a hardcoded boundary of 20 years, where childhood ages are transformed using a logarithmic function up to this boundary, while adult ages are untransformed. [9]. This biomarker generates very accurate predictions of chronological age, typically within a couple of years, but leaves many questions unanswered. Is there truly a change in the rate of epigenetic aging, from logarithm to linear trends, at 20 years? Does the linear fit of epigenetic age persist indefinitely, even for older individuals? Do non-linear trends in epigenetic aging vary across populations? As the biomarkers are species specific, they also do not allow one to directly address whether epigenetic aging trends vary across species. Moreover, recent studies [15, 3] question the validity of the time linearity assumption even in human adults, calling to a more global view of the process.

To address some of these questions, we have previously proposed a common framework by borrowing from the field of evolution. The universal pacemaker (UPM) of genome evolution was devised in the setting of molecular evolution in order to relax the time-linear evolution (i.e. rate constancy) imposed by the molecular clock hypothesis [20, 17], to account for correlation between rate changes in the genes of an evolving organism. The UPM is a statistical framework under which the relative evolutionary rates of all genes remain nearly constant whereas the absolute rates can change arbitrarily. In [19] we first proposed the adaptation of the UPM to the epigenetic setting, named the *epigenetic pacemaker* (EPM), and showed its application to simulation and small scale biological data. To the best of our knowledge, the EPM is the first model based framework for epigenetic aging, where the rates of change with time of individual CpG sites are parametrized, along with the epigenetic age of the individual. In [18] we devised a fast, conditional expectation maximization (CEM) algorithm that is capable of processing inputs of several thousands of sites and individuals.

In this work we set out to address some of the questions mentioned above, regarding the non-linear trends of epigenetic aging across populations. First we show that the EPM, which lacks any predefined regimes of age intervals, can be used to model and identify epigenetic aging trends over the entire lifespan of a population. We first apply our approach to a synthetic model simulating a non-linear aging process and show that our framework is capable of capturing the trend built into this model. Next we apply our model to publicly available sets of DNA methylation collected across broad age ranges and diverse tissues. Our results suggest unambiguously that a logarithmic trend across the entire lifespan is a better description of epigenetic aging than linear or polynomial trends.

## 2 Methods

### 2.1 The Evolutionary Models

Our basic objects are a set of *m individuals* and *n methylation sites* in a genome (or simply sites). Each individual has an age, forming the set *t* of *time periods* {*t*_*j*_} corresponding to each individual *j*’s age. Henceforth we will interchangeably refer to individuals with their age. Each individual has a set of sites *s*_*i*_ undergoing methylation changes at some *characteristic rate r*_*i*_. Each site *s*_*i*_ starts at some *methylation start level* 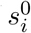. All individuals have all the sites *s*_*i*_. As *r*_*i*_ and 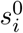 are characteristic of the site *s*_*i*_, by the model they are the same at all individuals. The latter fact, links the same sites across different individuals, but also within individuals by the fact that sites generally maintain the same characteristic rates across the whole population. Henceforth, we will index sites with *i* and individuals with *j*.

Now, let *s*_*i,j*_ measure the methylation level at site *s*_*i*_ in individual *j* after time (i.e. age) *t*_*j*_. Hence, under the *molecular clock* model (i.e. when rate is constant over time), we expect: 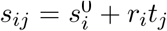. However, in reality we have a *noise effect ε*_*i,j*_ that is added and therefore the *observed value* 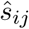 is 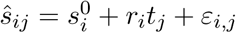.

Our goal is to find, given the input matrix 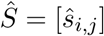, the maximum likelihood (ML) values for the variables *r*_*i*_ and 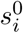 for 1 *≤ i ≤ n*. For this purpose, we assume a statistical model for *ε*_*i,j*_ by assuming that it is normally distributed, *ε*_*i,j*_ *∼ N* (0, *σ*^2^). In [19] we showed that minimizing the following function, denoted *RSS*, is equivalent to maximizing the model’s likelihood

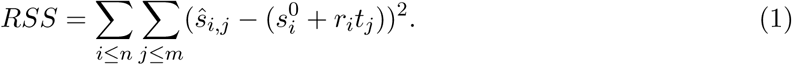

We also showed that there is an efficient and precise linear algebra solution to this problem, that we describe in more detail in the supplementary text.

In contrast to the MC, under the EPM model, sites may arbitrarily and independently of their counterparts in other individuals, change their rate at any point in life. However, when this happens, all sites of that individual change their rate proportionally such that the ratio *r*_*i*_*/r*_*i′*_ is constant between any two sites *i, i*′ at any individual *j* and at all times. In [19] we showed that this is equivalent to extending individual *j*’s age by the same proportion of the rate change. The new age is denoted as the *epigenetic age*. Therefore here we do not just use the given chronological age but estimate the age of each individual. Hence under the EPM we must find the optimal values of 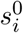, *r*_*i*_, and *t*_*j*_ (where *t*_*j*_ represents a weighted average of the rate changes an individual has undergone through life). The solution to this optimization problem is described in detail in our previous publications [19, 18]. We note that the deviation between the chronological age and the estimated epigenetic age is an age difference which, when positive, is denoted as age acceleration, and deceleration - otherwise.

To compare between the two models - MC and EPM, we note the following. The MC model is restricted to linearity with time by estimating a constant rate of methylation at each site, and using the given chronological age of each individual. The competing, relaxed, model (EPM) has no such restriction, and we estimate an “epigenetic” age for each individual. By definition, the ML solution under the relaxed model cannot be worse than the constrained model. For that specific case, when one hypothesis generalizes another, there is a special test, the likelihood ratio test (LRT), in which the specific hypothesis serves as the null hypothesis and the goal is to reject it in favor of the alternative one. In the supplementary text we provide a more detailed explanation of this test and its application in our case.

### 2.2 Selecting Informative Methylation Loci

DNA methylation platforms usually measure several hundreds of thousands of sites. It has been observed that many of these sites are invariant and do not change with age. It is desireable to retrictte analysis only to the most informative sites. Nevertheless, among the sites that do change it is necessary to set a criterion for site selection as it is inefficient to analyze all of the sites. There are several alternatives that we now describe.

The first and most basic and intuitive criterion for site selection is site variance - simply choose the sites that exhibit the largest variability. Figure 1(L) depicts the resulted analysis based on this criterion. This criterion is crude in the sense that it entirely ignores the relationship between time (age) and methylation. Therefore the next criterion to be examined is the covariance between age and methylation status at the site. The covariance metric selects sites that have a large change in methylation with age. We note that this criterion will not necessarily yield a significant linear fit between age and methylation status, the sites may still have a significant scatter, as is shown in Figure 1(M).

**Figure 1:**
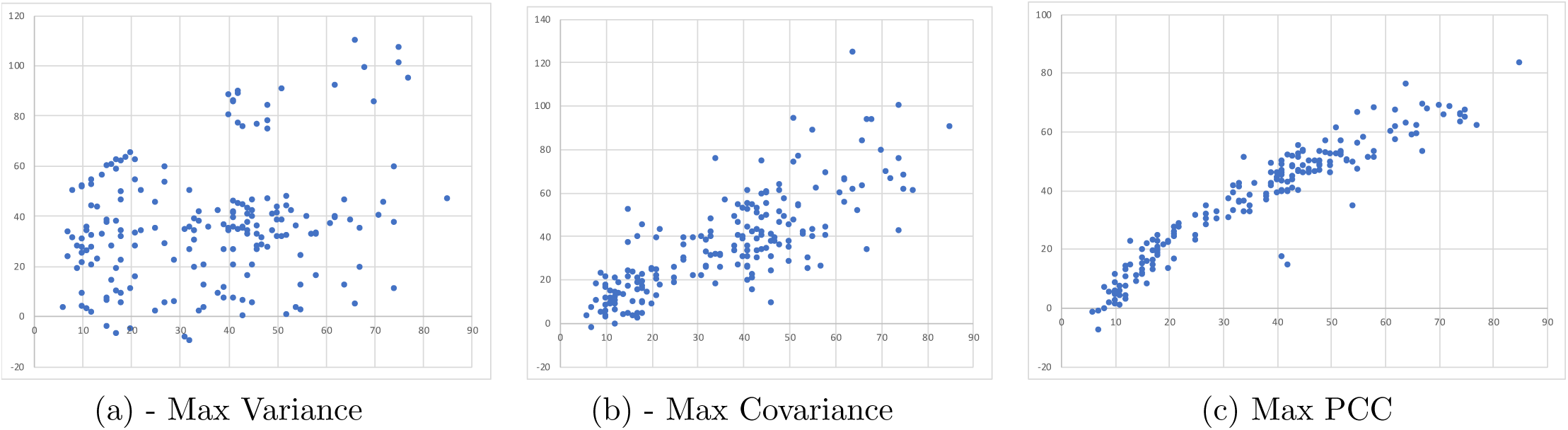
Site Selection Criterion. Scatter plots of inferred epigenetic age (e-age, y-axis) as a function of the chronological age (c-age, x-axis) as a result of applying the EPM algorithm to blood samples from data set GSE60132 (see more details in the Results sec.). Each point represents an individual. 1000 best sites were selected by the following three criteria. **Left:** Sites are selected based on their variance, regardless of correlation to age. **Middle:** Sites are selected based on their covariance with age. **Right:** Sites are selected by the (absolute) Pearson correlation coefficient.

Therefore the third criterion is the (absolute) Pearson correlation coefficient (PCC) defined as *ρ*_*X,Y*_ = *Cov*(*X, Y*)*/σ*_*X*_ */σ*_*y*_. In contrast to the covariance, PCC selects sites that have a tight fit to a linear relationship between methylation and age, although some of them show small changes in methylation across the range. Figure 1(R) shows the results based on sites selected by the PCC criterion. It is noticeable that using this criterion a much tighter relationship between epigenetic and chronological age is obtained. Hence we use PCC to select the sites to be modeled for all our data sets as it provided the clearest trends between epigenetic and chronological age.

### 2.3 Determining the trend line of epigenetic age

To determine the trend line between epigenetic and chronological age we employed both coarse and fine grained procedures based on the individual e-age inferred initially by the pacemaker criterion. Recall though that a first stage test is whether the pacemaker criterion, stating that rates and starting states are statistically correlated across individuals, and sites at any individual are also correlated, holds. This, is done by comparing to the molecular clock to the pacemaker model, as was described in the model description. Indeed, in the supplementary text we show the results of this test, along with the specific values obtained. The values depict that the pacemaker alternative is always superior with p-value smaller than 10^–6^.

We start by describing the two-stage procedure for determining the type of trend in the population. The mutual independence of the stages, along with lack of any prior assumption of any trend in the population, guarantees that the trend inferred is objective and unbiased. The EPM procedure is applied to the data in order to find optimal values for rates, starting states, and epigenetic age for all sites and individuals. Note that except for that pacemaker principle, that enforces uniformity of site rate and starting state across all individuals, there is no mechanism imposing correlation between any two individuals. Therefore, the EPM assigns every individual the optimal epigenetic age.

Once the EPM procedure is done, each individual is assigned its own epigenetic age. At the second stage, we seek a function that best fits the relationship between epigenetic and chronological age across the entire age range. Our prime criterion for goodness of fit for a trend of this relationship between epigenetic and chronological age, is the *R*^2^ coefficient from the trend line. In all our data sets we parametrized three functional forms for the trend line: linear, quadratic and exponential, and we used Excel to fit the best coefficients for each type. In the Results section we provide a more detailed description on this process.

We now describe a second approach that we devised and utilized. In age ranges where the trend is not conspicuous, that is, near linear, such that it cannot be distinguished convincingly using the above functional forms, we note the following. For a person *j* with age *t*_*j*_ and inferred e-age *p*_*j*_ we define *ρ*_*j*_ = *t*_*j*_*/p*_*j*_. Now, assume the increase in e-age is decreasing with time, then we observe that if we order the *ρ*_*j*_s by increasing *t*_*j*_, we obtain a monotonic increasing series (of *ρ*_*j*_). We denote that series [*ρ*_*j*_]. However, due to biological and statistical noise we never expect to find strict monotonicity at [*ρ*_*j*_] and we are bound to test only a *trend* of monotonicity. Now, we note that by the definition of *ρ*_*j*_ also the variance of *ρ*_*j*_ is changing in time. However, we note the following. For any two indices *j*_1_ and *j*_2_ such that *j*_1_ *< j*_2_, if [*ρ*_*j*_] is monotonically decreasing, then *ρ*_*j*1_ < *ρ*_*j*2_. Moreover, suppose we randomize the order of [*ρ*_*j*_]. Then for any two indices *j*_1_ and *j*_2_ the probability ℙ [*ρ*_*j*1_ *> ρ*_*j*2_] = 1*/*2 and hence the expected number of *j*_1_ *< j*_2_ such that 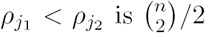. Now, since the variables *ρ*_*j*_ might be dependent, we cannot use standard bounds on deviations to calculate the probability of seeing that many pairs *j*_1_ *< j*_2_ such that *ρ*_*j*1_ *< ρ*_*j*2_ by chance. This forces the use of a non parametric test of the hypothesis. For this purpose we can use the Mann-Kendall test for monotonic trend [13, 12, 6]. According to this method, given a random vector *v*, all pairs of indices (*i, j*) such that *i < j* are checked whether *v*_*i*_ *< v*_*j*_. Let *p* be the number of pairs (*i, j*) for *i < j*, such that *v*_*i*_ *< v*_*j*_ and let *q* be the number of such pairs such that *v*_*i*_ *> v*_*j*_. Now let *S* = *p – q*, representing first the direction of trend with *S >* 0 when the series is increasing, and vice versa for *S <* 0. However, *S* also indicates on the intensity of the trend, and we note that under *h*_0_ (no monotonicity), we have *E*(*S*) = 0. Now we also need to compute the variance of *S*, 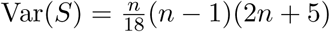. The statistic *z* defined 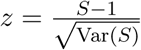 follows approximately the standard normal distribution, hence allowing us to obtain conveniently a *p*-value for the trend indicated by *S*.

## 3 Results

### 3.1 Identifying Trends in a Cohort

Methylation trends of the relationship between epigenetic and chronological time in a cohort provide useful information of how a group, as opposed to an individual, ages epigeneticlly with time. The Horvath model [9] has a rigid assumption of linearity of e-age in time for adults by using a linear combination of an individual’s methylation states of several hundreds of sites. For kids (age less than 20), the model corrects for non linearity using a logarithmic, yet fixed, function. The EPM model has no such assumption and therefore has the freedom to assign each individual its own e-age, as long as it complies with the EPM universality law, that is, that this age affects all the individual’s sites. We now demonstrate on synthetic data, the ability of our procedure to infer correct times (e-ages) and in particular trends throughout a whole population. For this purpose we have devised the following age related function that appears to encompass the characteristics of e-aging as they emerge from existing knowledge, in particular by the Horvath model:

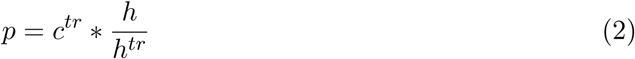

where *p* holds the e-age, *c* is the chronological age (c-age) of a person, *h* is some upper limit on a person’s age, and *tr* is a *trend* parameter to the function. The trend function has few desired characteristics. First, it satisfies a monotonic decrease in rate through time (c-age) and that decrease is proportional to the trend parameter. Also, at c-age that equals the upper limit *h*, the epigenetic and chronological ages coincide: *c* = *p*. Finally, for *tr* = 1 the trend function is linear with *p* = *c* for every *c*, as 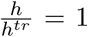. Figure 2(L) illustrates pictorially the behaviour of the trend function for several values of trend *tr* = 1, 0.8, 0.5, 0.1 and for upper limit age *h* = 100. Indeed we see that all trend lines depart from the origin and converge towards the point *c* = *h*. We also see that the larger *tr* is (with maximum *tr* = 1), the more straight the trend line is, and in particular, for *tr* = 1 a straight line with slope 1 is exhibited.

**Figure 2:**
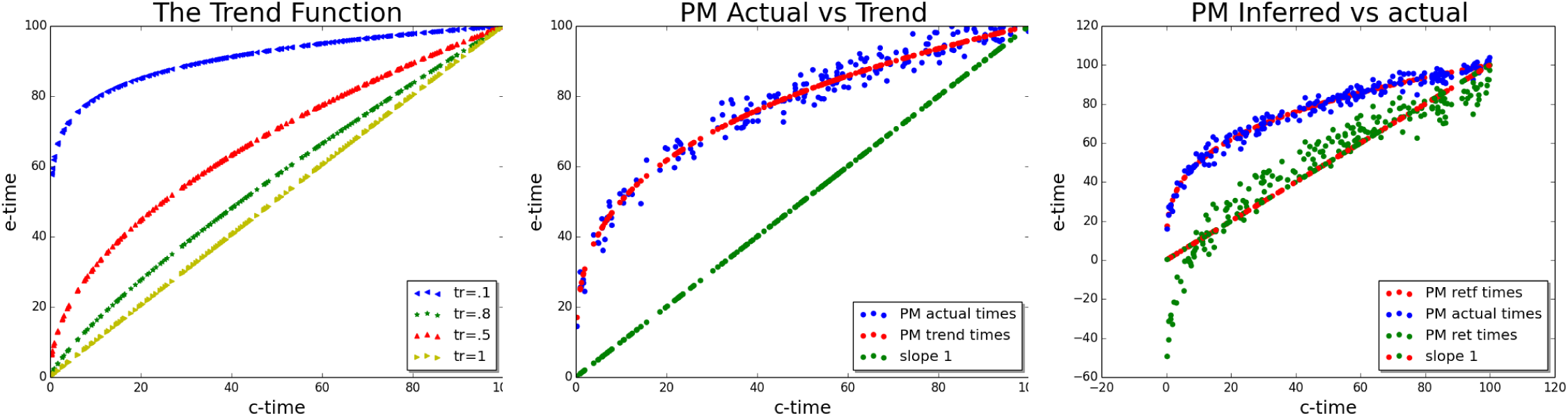
The trend function. **Left:** *The trend function* - Trend lines for four *tr* values *tr* = 0.1, 0.5, 0.8, 1 in blue, red, green, and olive green colors respectively. **Middle:** *Simulated actual noisy PM* - Actual noisy e-ages (blue dots) values around the trend line (red) with specific *tr* = 0.5 and *σ*_*p*_ = .8. The (green) 45*°* line represents the c-age of each individual. **right:** *The values inferred by the EPM-CEM algorithm* - green dots represent the inferred e-age by the algorithm. It should be compared to the real e-age (blue). While there is a gap, linear with time, between actual and inferred e-ages, the trend is captured.

In order to simulate realistic e-ages, we allow for each individual, some stochastic deviation of her/his e-age from the (or population’s) trend line, and that deviation depends on some variance *σ*_*p*_. To show that, we set a specific *tr* and *σ*_*p*_. Figure 2(M) shows simulated e-ages around the trend function with specific trend *tr* = 0.5 and *σ*_*p*_ = .8. For illustration, the straight 45° line, representing the c-age, appears in green the figure.

Finally, for every such e-age produced, our simulation procedure generates the methylation status 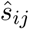 for every site *i* and and individual *j*, according to the model: 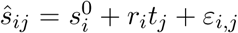. Figure 2(R) shows the result of applying the EPM-CEM procedure to such synthetic data. Green dots represent the inferred e-age by the algorithm and should be compared to the real (model) e-age (blue, same as in the middle box). We can see that EPM-CEM is capable of capturing the trend imposed by the simulation however it lags below the trend line by a gap that is linearly (inversely) correlated with age (red-green dots). This gap, returned by the procedure is due to the degeneracy of the likelihood surface that allows for multiple points in the surface to attain the same likelihood and in particular the maximum likelihood.

### 3.2 Analysis of Human Data

We have shown above the ability of our technique to identify the correct trend in aging in simulated data, and we now move to analyze human methylation data from various types of data sets. The general procedure we have taken to identify and assess a trend is as follows. We applied the EPM approach to the real methylation data, all taken from the Gene Expression Omnibus (GEO) repository, using the procedure in the simulation section above 3.1. The EPM, allows us to determine whether the MC hypothesis is rejected by the pacemaker, and also infers for each individual its epigenetic age. We remark that for all the real datasets we analyzed here, the pacemaker hypothesis was found superior to the linear approach with p-values always smaller than 10^−6^. As these information is not essential to the main subject of this study, it appears in details in the supplementary text.

Similarly to the simulation study, in a subsequent stage to the EPM, we plot for each individual its two ages. Here however, as these points were not synthetically generated by a function, we attempt to fit a trend function best describing these points. We focus on two families of functions - linear and logarithmic, as they are very general with a single explaining variable. These families are indeed the most common for trend approximation. However, to obtain additional insights on the functional form of the trend, we also accompany the logarithmic and the linear approximations with a quadratic best approximation line. In the results below, we demonstrate how we exploit the added information provided by the quadratic approximation. We note that any linear line is a special case of a quadratic family, simply with quadratic coefficient equals zero, and hence by definition its fit is always inferior to the quadratic approximation.

Our analysis is divided to age range based analysis, and also to tissue based analysis. All data sets required a preprocessing step of selecting the most informative (1000) sites, and based on our conclusions above, we used the Pearson correlation coefficient (PCC) criterion.

#### 3.2.1 Epigenetic Aging in Adults

Our first data set is from GSE40279, consisting of 656 blood samples from adults [7]. Their ages range from 19 years to 101. The results are depicted in Figure 3. In the top two graphs we show scatter plots with the three trend lines where the one with the best fit appears in the left and the two suboptimal ones in the right scatter plot. For each trend line, we also depict its exact formula and the *R*^2^. It is quite evident that the linear trend is superior here to the other two, with negligible increase in *R*^2^ for the quadratic trend. Therefore, in order to check if there is still a trend of non linearity, we applied the Mann-Kendall test as described in the Methods section. The lower part of Figure 3 shows the values of *ρ*_*i*_ ordered by chronological age. We set to test if there is an increasing trend in this series. The value obtained for *S* was 1619 implying that we have an increasing trend in *ρ* and therefore epigenetic aging rate is decreasing in time. However, under this size of data, we also have a huge variance with *V ar*(*S*) = 31438253.33 yielding *z*-score of 0.28856 which is not significant.

**Figure 3:**
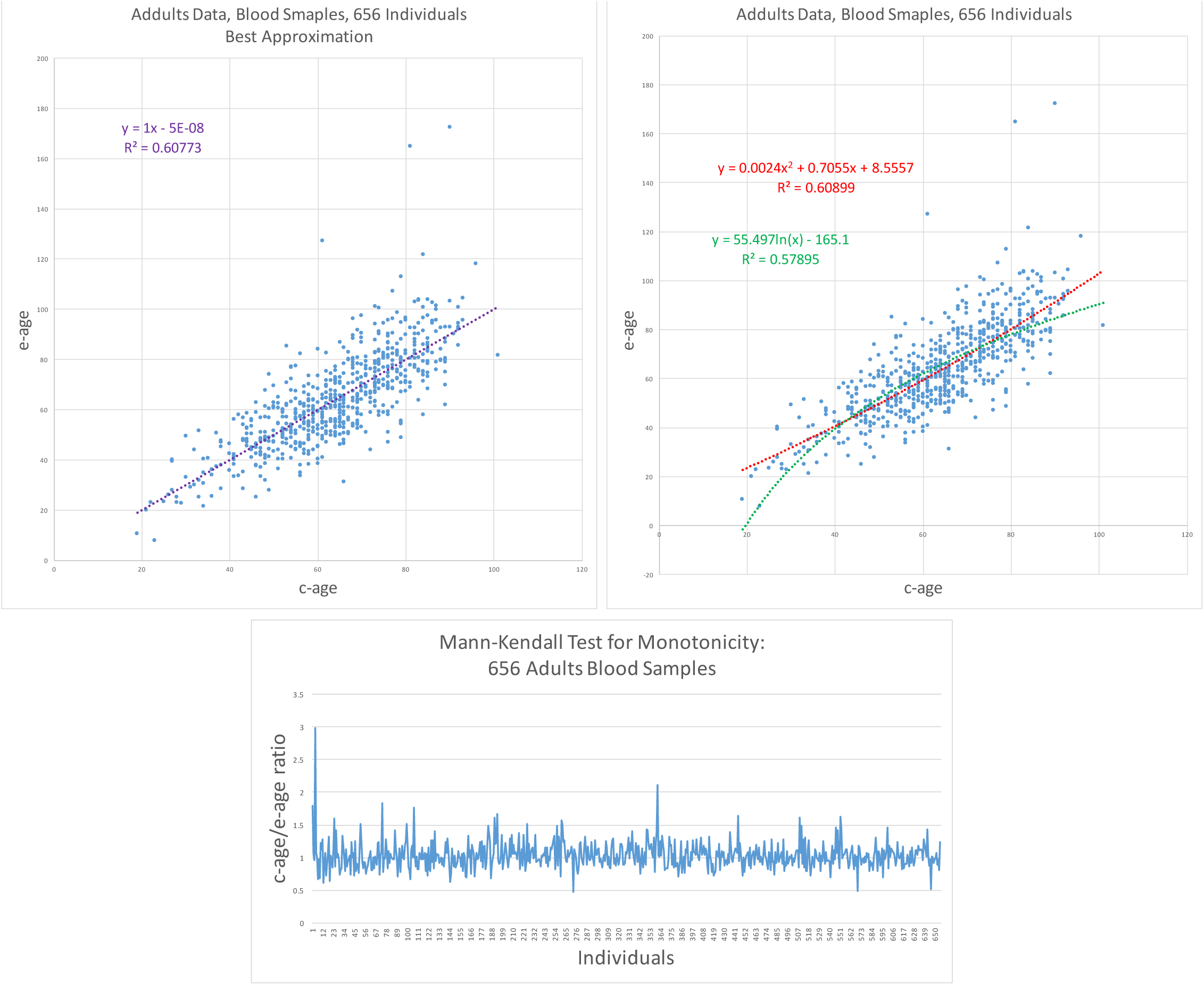
GSE40279 - Human Blood Data Results I. (**Top**) E-age vs C-age in adults. Age is plotted in years. The left graph shows the best approximation to the data. The linear line is slightly and insignificantly inferior to the quadratic approximation and therefore is the best fit. (**Bottom**) Mann-Kendall Test for monotonicity Trend: c-age vs e-age ratio ordered from left to right according to c-age. If rate of aging is decreasing, we expect to see monotonic increase in the function. Indeed the function is increasing but not in a significant manner.

Our second data set is the GSE87571, also from human blood taken from 366 individuals of ages from 14 to 94 years old. The results of applying the EPM-ECM to this data are depicted in Figure 4. The top two graphs show the scatter plot of e-age versus c-age with the three trend types - linear, quadratic, and logarithmic. The linear trend line in the left is only slightly inferior to the quadratic with *R*^2^ = 0.612 versus *R*^2^ = 0.613 and therefore we consider it to be the best approximation. Nevertheless, the advantage in *R*^2^ of the linear trend over the logarithmic trend is only .5% and indeed here we already see a bend in the points corresponding to younger ages. In general, the entire collection of points here allude to a concave shape, i.e. a decreasing function, as can be seen by the negative coefficient (-0.0045) of the quadratic term in the quadratic trend line. This should be contrasted to the convex trend of the quadratic trend in the previous case (data set GSE40279 - Human Blood Data, Fig 3) where the first coefficient equals 0.0024.

**Figure 4:**
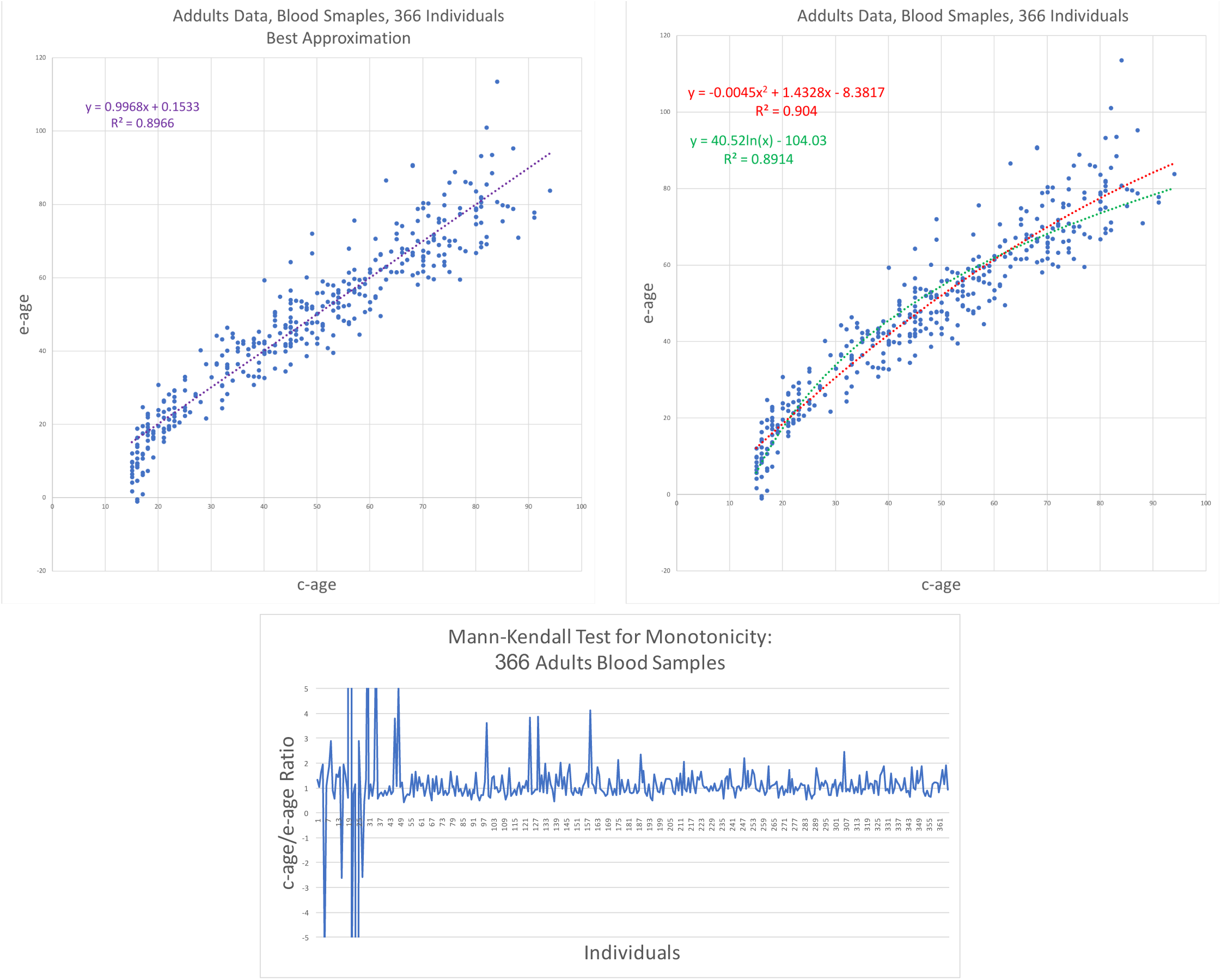
GSE87571 - Human Blood Data Results II. (**Top**) E-age vs C-age in adults. Age is plotted in years. The left graph shows the best approximation to the data. The linear line is slightly and insignificantly inferior to the quadratic approximation and therefore is the best fit. (**Bottom**) Mann-Kendall Test for monotonicity Trend: c-age vs e-age ratio ordered from left to right according to c-age. If rate of epigenetic aging is decreasing with time, we expect to see a monotonic increase in the 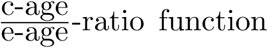. Indeed the function is significantly increasing.

To verify this decreasing trend, as in the previous data set, we apply the test of monotonicity - the Mann-Kendall Test - to this data. The value obtained for *S* was 4225 implying that we have an increasing trend in *ρ* and therefore epigenetic aging is decreasing in time also for this data set. The variance here is *V ar*(*S*) = 5469768 yielding a *z*-score of 1.806 and a p-value smaller than 0.04 and is therefore significant.

#### 3.2.2 Epigenetic Aging in Children

After analyzing the data collected from adults, we turned to analyze data from children. We analyze the GSE36064 data set of blood samples taken from 78 children of ages ranging from one year to 16. The ages here, as well as in the figure describing it, are represented in months. The results are shown in Figure 5. Here, the linear trend is clearly inferior to the quadratic and the logarithmic trends. We find that the logarithmic trend is the best approximation and show it in the left side of Figure 5.

**Figure 5:**
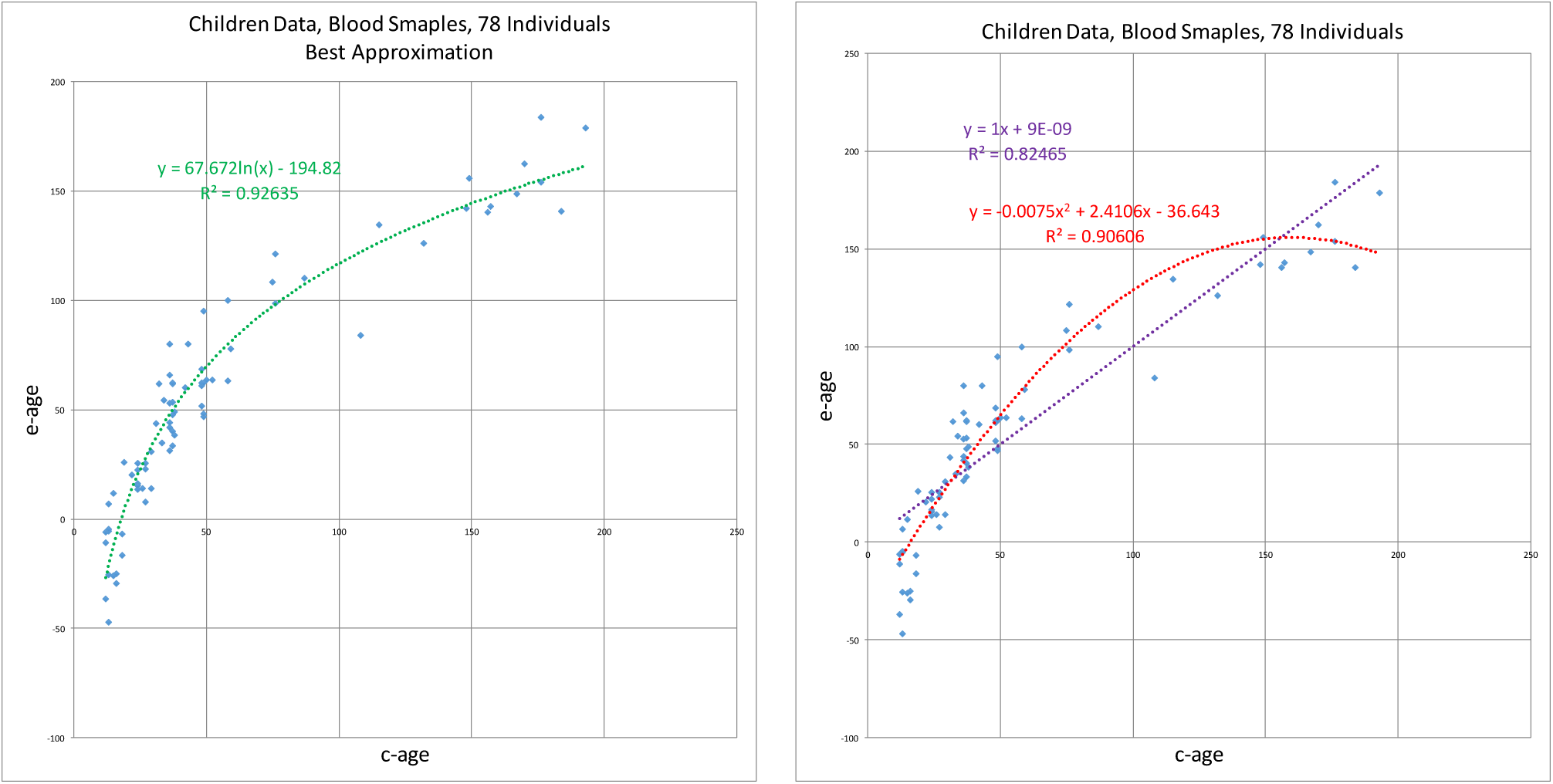
GSE36064 - Children Blood Data Results. E-age vs C-age in young humans. Age is plotted in months. The left graph shows the best approximation to the data. The logarithmic approximation provides the best explanation.

#### 3.2.3 Combined Age Analysis

In the previous data sets we restricted the analysis to specific age ranges, such as children or adults. In the next two data sets we analyze blood samples from individuals with age ranges from childhood to old age. The first data set - GSE60132 - was taken from peripheral blood samples of 192 individuals of Northern European ancestry [1]. Ages range from 6 to 85 years. The results are shown in Figure 6. As can be noticed, the logarithmic trend line provides better *R*^2^ than the linear trend line, 0.9125 versus 0.8998, and even better than the quadratic trend line (*R*^2^ = 0.9119). The concavity of the spread of the points is fairly noticeable and this is confirmed by the negative first coefficient of the quadratic trend function - −0.0078.

**Figure 6:**
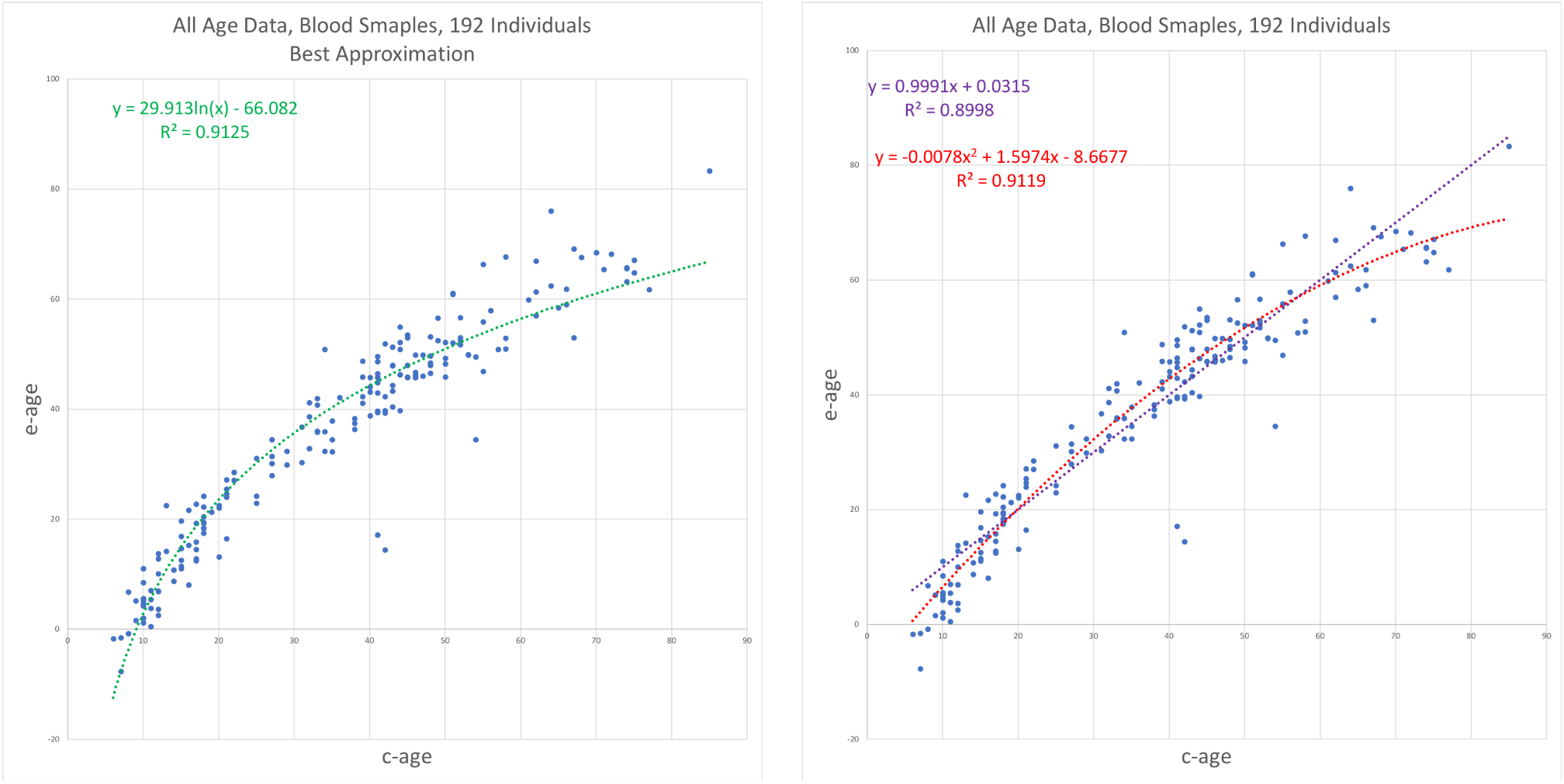
GSE60132 - Human, All Ages, Blood Data Results I. E-age vs C-age in wide age range. Age is plotted in years. The left graph shows the best trend line approximation to the data, which is the logarithmic trend function. At the right, the inferior trends - the quadratic and linear. The quadratic line is slightly and insignificantly inferior to the logarithmic approximation, buy also portrays a concave line due to negative first coefficient -0.0078.

The Next data set - GSE64495 - is also from blood samples of 113 individuals [22]. Here, while there is a scarcity of samples from the age range 12-35, the entire age range of the study begins at even younger ages than the previous data set: 2.3 years versus 6. Our results for this dataset are depicted in Figure 7. Here the advantage of the logarithmic trend line over the linear is the most significant among the adults containing data sets analyzed so far, *R*^2^ = 0.97534 versus *R*^2^ = 0.8664, and is even significant over the quadratic - *R*^2^ = .9035. The decrease in the rate is evident as well as the fit to the logarithmic trend line.

**Figure 7:**
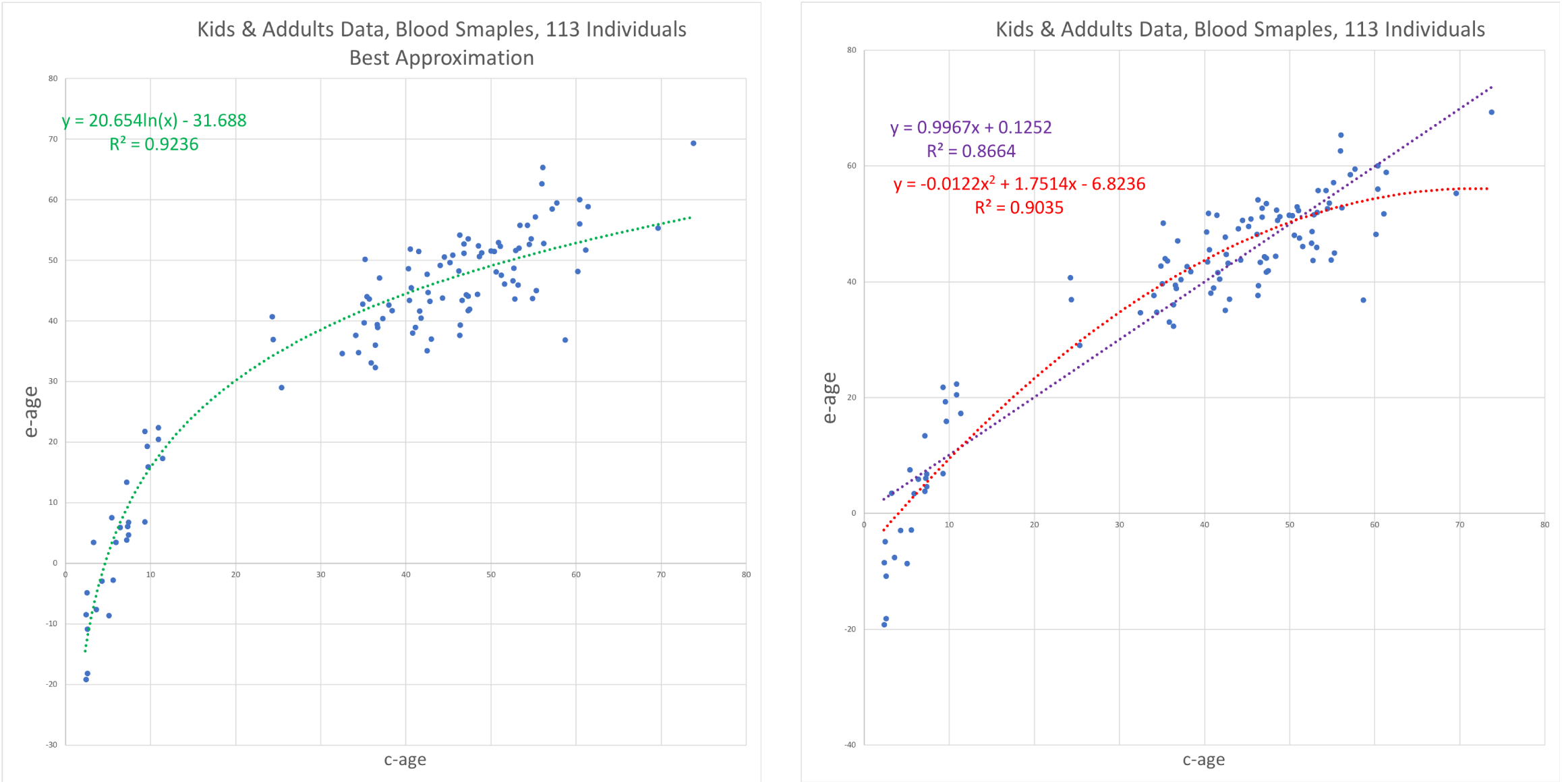
GSE64495 - Human, All Ages, Blood Data Results II. E-age vs C-age in kids and adults. Age is plotted in years. The left graph shows the best approximation to the data, obtained by the logarithmic trend line with *R*^2^ = 0.9236. On the right, the inferior trend lines, the linear line with *R*^2^ = 0.8664 and the quadratic line with *R*^2^ = 0.9035.

### 3.3 Brain Development and Aging

Our last data set is from GSE74193, consisting of 675 samples from brain tissues from before birth to old age [10]. The advantage of this data set is two-fold. First, the broad range of ages - from half a year before birth to 85 years, which represents a broader age range than that found in the the previous data sets, and allows us to track epigenetic aging across the entire span of life, starting from before birth. Second, all the samples from the previously analyzed data sets came from blood. This data set, from brain tissues, allows us to contrast our results from blood tissues to another tissue type. Our results are depicted in Figure 8. The logarithmic approximation, appears on the left graph, not only provides a significantly better fit to the data, with *R*^2^ = 0.97534 versus *R*^2^ = 0.86448 and *R*^2^ = 0.70708 for the quadratic trend line and the linear trend line respectively. Moreover, the high *R*^2^ provides an almost perfect fit to the data.

**Figure 8:**
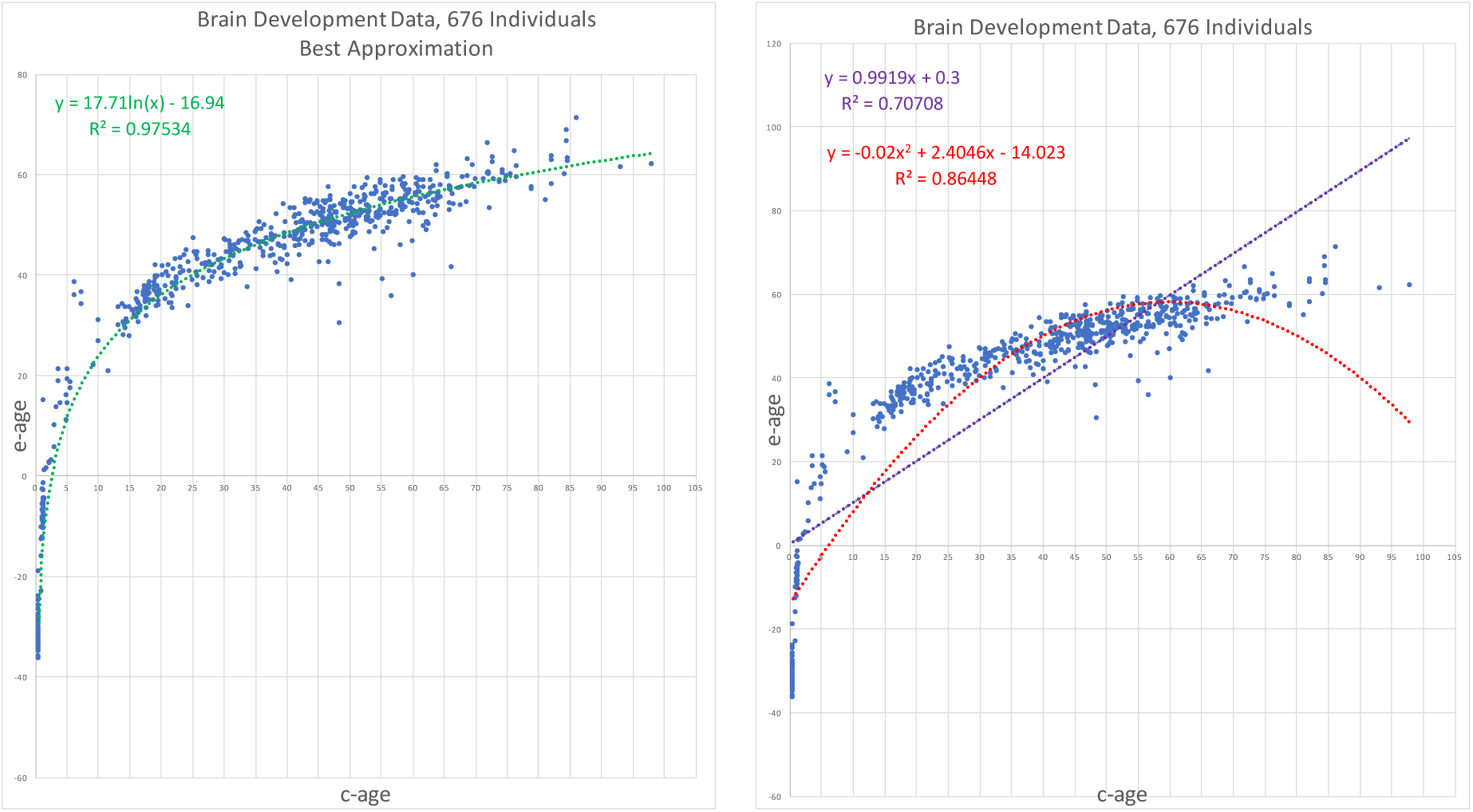
GSE74193 -Brain development Data Results. E-age vs C-age in young humans. Age is plotted in years. The left graph shows the best approximation to the data. The logarithmic approximation provides the best explanation.

## 4 Discussion

During the past few years several studies have shown that DNA methylation patterns continue to change as individuals age. These observations have been leveraged to construct epigenetic clocks that predict the age of an individual based on their methylation profile. While these tools have proven to be very useful for aging studies, they are based on a priori assumptions about the relationships between epigenetic and chronological age. For example, the Hannum age clock assumes that epigenetic age and chronological are linearly related, and using multivariate penalized regression identifies 71 CpG sites whose methylation values can be combined using a weighted sum to predict the actual age of the individual. The Horvath clock uses a more complex set of assumptions to derive its predictions of chronological age: it applies a logarithmic transformation to ages below 20 and a linear relationship for ages greater than 20. He then identifies 353 CpG sites using elastic net regression to very accurately predict the transformed age. These biomarkers have been widely used to study aging, as is evident by the hundreds of studies that have utilized them. However, because of their underlying assumptions, they are not ideal tools to infer trends in epigenetic aging rates across life spans in an unbiased fashion. However, correctly modeling potential nonlinearities in epigenetic aging is critical to advance the field and generate an even more robust understanding of epigenetic aging and its impact on human health and mortality.

To address this question, we have developed an unbiased approach to measure the trends in epigenetic aging within a cohort of individuals of varied ages. Our approach is distinct from the Hannum and Horvath epigenetic clocks in that it is not designed to optimally predict the age of an individual, but rather to model the non-linear trends in epigenetic aging over time without making any a priori assumptions about what these may be. The method is inspired by evolutionary models that attempt to model mutation rate changes over time. This lead to the development of our epigenetic universal pacemaker model, which we have presented in previous studies [19, 18].

To attempt to identify rates of epigenetic aging over the entire life span of humans we apply the epigenetic pacemaker to multiple datasets that measure DNA methylation in large cohorts of individuals of varying ages. In the first few cohorts the methylation is profiled from blood, while in the last cohort the methylation is measured in brain tissue. In all cases methylation is profiled using an Illumina microarray that measures the methylation across approximately 450,000 sites. While all cohorts sample individuals from early adulthood to old age, the second set also include samples from individuals from early childhood to adulthood, and in the last case of the brain study, also samples from fetuses obtained before birth.

By applying our epigenetic pacemaker model to this data we observe consistent and robust trends across these datasets. The first is that from early adulthood (around age 20) to old age (well into the 90s), DNA methylation changes in a roughly linear fashion, yet with a slight but significant tendency for rate decrease. However, in contrast to the adults, we find that DNA methylation changes are strongly nonlinear from late fetal stages to adolescents. In both the blood and brain datasets that measure these stages we observe that the epigenetic age inferred by our model is related to the chronological age by a logarithmic transformation across the entire span of life, from before birth to old age. Thus, DNA methylation changes are very rapid initially, and then gradually decrease with age. This implies that the rate of change of epigenetic ages (i.e. the slope of our trend line) is roughly the inverse of the chronological age. The fact that we consistently observed these trends across multiple datasets, and two different tissues, suggests that the logarithmic relationship between epigenetic and chronological age may be a universal property of human aging.

This universal logarithmic trend may help explain some interesting observations that have been reported in the literature regarding epigenetic aging. For example, one recent study found that the Horvath epigenetic clock “systematically underestimates ages in tissues from older people,” and that “a decrease in slope of the predicted ages were observed at approximately 60 years, indicating that some loci in the model may change differently with age, and that age acceleration measures will themselves be age-dependent”[3]. A second study also found that “epigenetic age increases at a slower rate than chronological age across the life course, especially in the oldest population”[15]. These results suggest that the underlying assumptions about the relationships between epigenetic age and chronological age impact the performance of the Hannum and Horvath epigenetic clocks, and that deviations are most notable in old age. We speculate that if the logarithmic epigenetic aging trend that we observe is in fact universal, that this could lead to improved biomarkers that show more robust performance at the extremes of the age distributions, leading to more accurate associations between epigenetic aging and human health and longevity.

Moreover, we believe that the observation that epigenetic aging is logarithmic over the entire life span opens up new avenues for epigenetic research in the future. What mechanisms lead to the gradual reduction in epigenetic rates from late fetal stages to centenarians? Is the logarithmic trend related to prior observations that epigenetic aging is a measure of epigenetic entropy? The answers to these questions will undoubtedly influence our understanding of human aging and longevity and will most likely apply to a broad range of organisms. By quantitatively demonstrating these trends in an unbiased fashion, we believe we have laid a solid foundation for the development of improved aging biomarkers and the investigation of the underlying mechanisms of epigenetic aging, and ultimately to the answers to these important questions that are fundamental to the biology of development and aging.

